# AVOCADO: Visualization of Workflow-Derived Data Provenance for Reproducible Biomedical Research

**DOI:** 10.1101/044164

**Authors:** Holger Stitz, Stefan Luger, Marc Streit, Nils Gehlenborg

## Abstract

A major challenge of data-driven biomedical research lies in the collection and representation of data provenance information to ensure reproducibility of findings. In order to communicate and reproduce multi-step analysis workflows executed on datasets that contain data for dozens or hundreds of samples, it is crucial to be able to visualize the provenance graph at different levels of aggregation. Most existing approaches are based on node-link diagrams, which do not scale to the complexity of typical data provenance graphs. In our proposed approach we reduce the complexity of the graph using hierarchical and motif-based aggregation. Based on user action and graph attributes a modular degree-of-interest (DoI) function is applied to expand parts of the graph that are relevant to the user. This interest-driven adaptive provenance visualization approach allows users to review and communicate complex multi-step analyses, which can be based on hundreds of files that are processed by numerous workflows. We integrate our approach into an analysis platform that captures extensive data provenance information and demonstrate its effectiveness by means of a biomedical usage scenario.

## 1 Introduction

Recent advances in biomedical research enable the rapid acquisition of data from biomedical samples for clinical and pre-clinical studies. The bioinformatics workflows employed to analyze such data incorporate many distinct steps and tools that often result in long workflows. Moreover, the complexity increases with repeated workflow execution and number of processed samples. In the long run, the analyses associated with a biomedical study become hard to maintain, compare, and reproduce. To address this issue, all parameter modifications and workflow executions need to be captured as provenance information. Workflows may also be modified or executed multiple times with different parameters, or use a different dataset as input—resulting in a complex data provenance graph. Most existing visualization approaches are based on node-link diagrams, which usually do not scale well to large provenance graphs of dozens to hundreds of nodes. Hence, a major challenge is to effectively visualize such graphs in order to allow analysts understand the dependencies between different files in a dataset.

In recent work by Ragan et al. [RESC15], provenance is characterized by the supported provenance *type* and *purpose*. According to their organizational framework for provenance, our approach operates on *data provenance* containing all executed workflows together with their parametrization as well as their in-and output files. The purpose of our visualization is to *recall* the analysis history, enabling analysts to better understand complex analyses, and to *present* the information to both colleagues who are involved in project and others that are not part of the team, such as the general public. *Replication* and *action recovery* (undo/redo), however, are not direct goals of our visualization as this is typically handled by the tools that capture and manage data provenance information.

The primary contribution of this paper is AVOCADO (Adaptive Visualization of Comprehensive Analytical Data Origins), an interactive provenance graph visualization approach that visually aggregates the data provenance graph by exploiting the inherent topological structure of the graph. Based on the aggregation, we then expand relevant parts of the graph interactively using a multi-attribute degree-of-interest (DoI) function. As a secondary contribution, we integrate the visualization approach into the *Refinery Platform* [Ref15a] that captures, manages, and allows users to operate on data provenance information of bioinformatics workflows. We demonstrate the effectiveness of our visualization by means of a usage scenario.

## 2 Background

In the past decade, biomedical research has transitioned from being primarily hypothesis-driven to a data-driven endeavor. This has been the result of the availability of large amounts of heterogeneous data from genomics studies, electronic health records, imaging data, and other modalities. To analyze such data, complex bioinformatics workflows are employed that usually operate on files that are run through a series of specialized tools. In particular the analysis of genomic data, e.g., from studies that aim to identify driving mutations in cancer or to pinpoint genetic variants that are causing rare diseases, several multi-step workflows are commonly employed for quality control, preprocessing and data normalization, identification of statistically significant differences between cases and controls, identification of correlations, and other higher-level analyses, e.g., to identify changes in gene expression levels associated with a particular genomic mutation.

The failure to reproduce the results of a large number of such studies has raised major concerns in the biomedical research community and triggered several efforts to address this issue [BE12, HG13, BI15, Buc15]. The reasons why such studies fail to reproduce are diverse and range from inadequate statistical power, publication of incomplete or wrong study protocols, to unavailability of experimental raw data [Kai15]. In particular the failure to record and share data provenance information for published data frequently prevent results of computational analyses from being reproduced. Even when such information is published, it can be extremely challenging to reproduce a study [GBLZ*15]. A number of bioinformatics data analysis systems have been developed in recent years that aim to track data provenance automatically and comprehensively (e.g., *Kepler* [ABJF06], *Taverna* [WHe13], *Galaxy* [GNT10], *VisTrails* [BCS*05]). However, they lack adequate visualization tools to review and communicate this provenance information, which severely limits its value for reproducibility purposes.

A comprehensive effort to facilitate collaborative and reproducible biomedical research is the open source *Refinery Platform*, which integrates data management and analysis. Refinery handles data at the file level and facilitates the execution of workflows on one or more input files in the *Galaxy* bioinformatics workbench [Gal15]. For each of these analyses, Refinery automatically tracks comprehensive data provenance, including workflow templates applied, workflow parameters, tool versions, input files, the user executing the analysis, and others. Every *analysis* consists of one or more *analysis input groups*, which correspond to the execution of a Galaxy *workflow* on a set of input files (see Figure 2). The raw data sets are imported into Refinery as Investigation-Study-Assay (ISA-Tab) files [RSBM*10], which provide meta data and information about the generated raw data. For example, if a user selects 10 files to be processed by a workflow that takes one input file and produces one output file per input file, then the corresponding analysis would have 10 inputs and 10 outputs and would consist of 10 analysis input groups. Every analysis uses only exactly one workflow template. Along with the meta data attributes that users can assign to files in Refinery, the data provenance information represents a richly annotated graph that contains all information necessary to reproduce the findings of a study performed with the help of the system.

**Figure 1:**
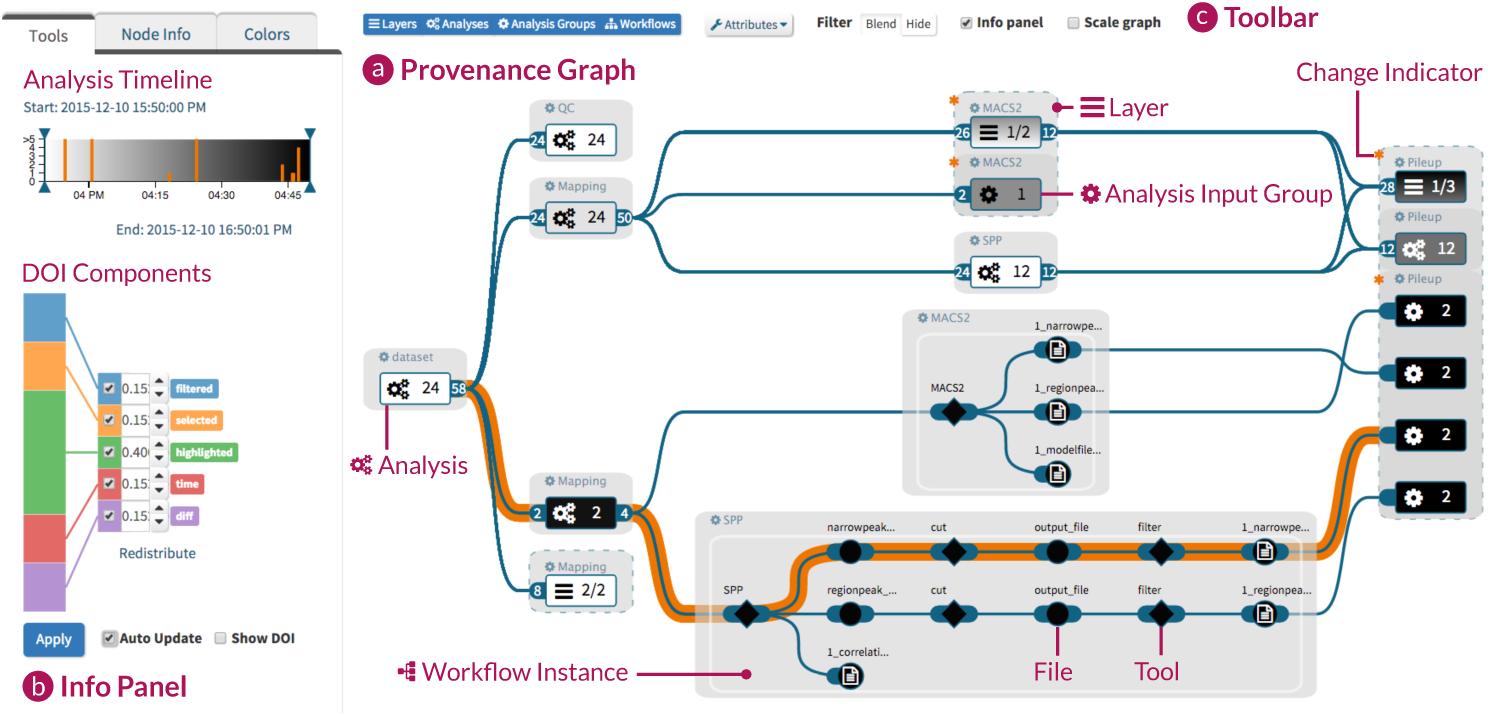
The provenance graph (a) is aggregated and filtered based on the selected analysis execution time and the weighted degree-of-interest (DoI) components (b). In the top center of the graph, two horizontally aligned workflows show a compound layer node, where the top node represents the layer itself while two workflows are extracted based on their specific DoI value exceeding a predefined threshold. The toolbar (c) allows users to switch between node type specific views (layer, analysis, analysis input group, workflow instance) and to change the attribute mapping onto nodes.

**Figure 2:**
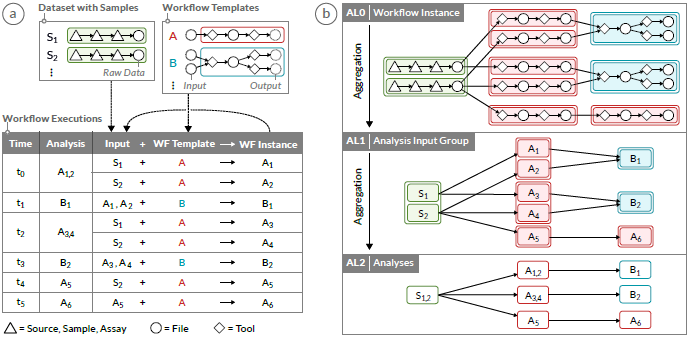
A workflow provenance graph consists of multiple workflow executions over time. (a) Each execution requires one or more input files and applies a workflow template, containing a sequence of tools and files. (b) The provenance graph can be visualized at multiple aggregation levels (AL) exploiting the hierarchical structure of the graph. AL0 shows the connected workflow instances that are first aggregated to analysis input groups at AL1 and further aggregated to analyses at AL2. The aggregation methods are explained in Section 5.1.

Data provenance graphs like those generated by Refinery and similar tools are DAGs. They contain different types of nodes, primarily file nodes and tool nodes. The latter represent the software that is used to process the files in a particular workflow. As illustrated in Figure 2, tools and files in workflow templates, analysis input groups, and analyses form a hierarchy that can be exploited to aggregate parts of the graph, as discussed in Section 5. Furthermore, data provenance graphs are often very broad, because each raw input file is run through the same set of workflows, either with the same or different tool parameter settings. The length of the path from a raw input file to a highly processed output file can include a dozen steps or more, making the graph not only broad but often also deep.

## 3 User Tasks

In close collaboration with bioinformatics experts and with the input from biomedical researchers, we refined Lee’s et al. task taxonomy for graph visualization [LPP*06] to identify a series of tasks that need to be supported by an effective data provenance graph visualization. The experts that we worked with are leading the development of the Refinery Platform and related projects. They include one of the authors of this publication. One of the authors who is a visualization expert worked very closely with the Refinery Platform team for more than 12 months.

**Task I: High-Level Overview** Users want to start the exploration by inspecting an aggregated version of the data provenance graph, to get an overview of which workflows were run and how often, in which configuration, and at which point in time. While further details should be hidden and summarized, an indication of the actual graph’s depth and breadth is desired.

**Task II: Attribute Encoding** Analyses are annotated with a series of attributes such as date and time of execution, in-and output files, and others. This information should be accessible through the provenance visualization.

**Task III: Drill-Down on Demand** Due to the space constraints, the graph should be presented as reduced as possible, but users still need to be able to view information at the highest level of detail. The visualization should enable drill-down operations into sub-graphs that are of current interest, while the rest of the graph should be kept in a compact representation as context.

**Task IV: Investigate Differences in Aggregates** The precondition for aggregating analyses is a common workflow template. However, tool parameters, input files, and the number of analysis input groups might be different, e.g., when an analysis was re-run with additional inputs or parameters. The data provenance visualization needs to provide users with the means to identify and investigate such differences.

**Task V: Investigate Causality** A crucial task in the exploration of data provenance graphs is to let users investigate the chain of files and transformations that contributed to a certain analysis result.

**Task VI: Search and Filter** Users should be able to focus on certain aspects of the graph, e.g., a specific workflow type, provenance data changes, or execution time range, by triggering filter and search actions.

Although these tasks were developed with the input of experts who are working in the biomedical domain, we expect that these tasks are also relevant to other application domains in which similar analysis pipelines are employed for data-driven research.

## 4 Related Work

Data provenance graphs are directed acyclic graphs (DAGs), which can lead to the conclusion that much of the graph visualization literature might be relevant. Two characteristics, however, make data provenance graphs special: (1) by design they include a hierarchy, and, (2) they have an inherent temporal aspect. We first summarize how suitable node-link diagrams and matrix representations—the fundamental graph visualization techniques—are for addressing the tasks introduced above. We then discuss different graph aggregation strategies and techniques suitable for visualizing dynamic graphs. We end our review with an overview of the state-of-the-art in provenance graph visualization.

Graph Representation The two fundamental techniques to visualize graphs are node-link diagrams and matrix representations. Which technique works better, depends on the graph type, graph size, and user tasks. Visualizing a graph as a matrix is well suited for attribute-based tasks performed on weighted edges, where each edge has an associated value [SS06]. However, for path-related tasks, such as following a path to address causality tasks (Task V), node-link diagrams are more effective. Therefore, node-link representations are better suited to the characteristics of data provenance graphs and for fulfilling the user tasks defined in Section 3.

**Graph Aggregation Strategies** The scalability of node-link diagrams is a well studied area of research with a large body of existing work. To improve the scalability for visualizing provenance graphs, we can utilize the hierarchy for aggregation [NJ04, EF10]—allowing users to explore the graph using drill-down (Task III) and roll-up (Task I) operations. In combination with semantic zooming [PF93], the information shown can be adjusted to various levels of detail, avoiding visual clutter. Further visualization techniques for group structures in graphs, as they appear through aggregation, were surveyed recently [VBW15]. An alternative aggregation method is motif discovery and compression [MSOI*02, AS06] which reduces the visual complexity through topology-based aggregation while preserving the basic structure of the graph [MRSS*13]. Instead of introducing new aggregations, large graphs can also be decomposed in multiple hierarchies [SGKS15] and visualized separately. Inlays shown on demand reintroduce the lost relationship between the hierarchies.

A different approach is to distort the visual space, as typically done in lens-based approaches [TAHS06]. Furnas [Fur86] visualized the nodes with different level of detail, determining the degree-of-interest (DoI) based on the selected node. The *DOITree* approach by Heer et al. [HC04] applies a multi-focal version of that DoI function to visualize tree structures more effectively. Van Ham and Perer [vHP09] extended the DoI approach to expand parts of a large static graph showing the context, preserving the overall graph structure around a selected node. Abello et al. [AHSS14] presented a DoI function that is divided into multiple components to investigate large dynamic networks. Vehlow et al. [VKB*15] use a combination of continuous and/or discrete DoI functions to filter dense biological networks and compare the extracted subnetworks subsequently. In this paper we propose to combine hierarchical and motif-based aggregation with a user-driven DoI to increase the scalability for exploration of large data provenance graphs.

**Dynamic Graph Visualization** In visualization, temporal aspects of data are particularly challenging because of the unique characteristics of time, such as the presence of hierarchical levels of granularity with irregular divisions, the occurrence of cyclic patterns, or the fact that time cannot be perceived by humans directly [AMST11] Researchers studied the temporal aspect also in the context of graphs [KKC14, HSS15]. When dealing with data provenance graphs, the user wants to investigate the differences between two or more analyses executed at different time points (Task IV). These differences can be visualized either by mapping time to time (*animation*) or time to position (*juxtaposition* and *superimposition*) [BBDW14]. Archambault [Arc09] uses superimposition of different snapshots in combination with hierarchical aggregation of adjacent nodes and path-preserving coarsening. Similarly, a recent approach by van den Elzen et al. [vdEHBvW15] allows users to explore the evolution of networks by reducing snapshots of the dynamic graph to points, forming a separate derived graph as an abstraction layer. These approaches visualize differences of large graphs well, however, they are not suitable for the data provenance graph problem, as they do not support path-related causality tasks (Task V).

**Provenance Graph Visualization** Over the past years, workflow and provenance management systems (e.g., *VisTrails* [BCS*05]) have become more effective at capturing and storing provenance information. However, these systems provide no or only a basic visual representation of this information. For example, *Synapse* [OEY*13] tracks data provenance information to ensure reproducibility in cancer genomics and other biomedical research and visualizes the data provenance graph as node-link diagram. However, this approach does not scale to large graphs, due to heavy usage of labels and icons, lack of visual glyph encoding, and missing aggregation techniques, which is in conflict with Task I. In contrast, *Provenance Map Orbiter* [MS11] uses aggregation techniques that compress the provenance graph to a high-level overview. Additionally, it uses semantic zoom and supports drill-down to show details on demand. However, the graph layout does not adapt well to dynamic aggregation, resulting in additional edge crossings—hampering Task V.

Overall, none of the discussed solutions is able to address all of the tasks formulated above. The challenge is therefore to design an effective combination of existing techniques and strategies for visualizing large data provenance graphs with hundreds of nodes.

## 5 AVOCADO Visualization Concept

In AVOCADO we reduce the complexity of the data provenance graph through a combination of graph aggregation and expansion strategies. Parts of the aggregated graph that are relevant to the user are expanded on demand by applying a modular degree-of-interest (DoI) function. This interest-driven adaptive approach allows us to handle complex multi-step analyses that can be based on hundreds of files processed by multiple workflows (see Section 2). Figure 1 illustrates our visualization using a data provenance graph with 13 analyses containing 927 nodes from different workflows.

### 5.1 Graph Aggregation Strategies

The first part of our approach reduces the data provenance graph through a combination of hierarchical and motif-based aggregation chosen to provide a meaningful overview that preserves the overall graph structure (Task I).

**Hierarchical Aggregation** Hierarchical aggregation reduces the size of a dataset by grouping related data items into aggregates based on the result of a recursive clustering operation. In our case we make use of the inherent hierarchy contained in the data provenance graph. We aggregate *workflow instances* (AL0) into *analysis input groups* (AL1) and further into *analyses* (AL2) (see Figure 2(b)). Analyses contain all analysis input groups that share the same workflow execution time. In AVOCADO we render all aggregation levels from top (AL2) to bottom (AL0) with the result that analyses, analysis input groups and workflow instances are stacked on top of each other. Each aggregation level is traversed in a breadth-first approach, placing the nodes in a column-based layout.

**Motif-Based Aggregation** A motif compresses a graph or parts of it while preserving the basic graph structure. Traditional motif discovery algorithms search for all permutations of a fixed amount of nodes.In AVOCADO motifs visualize the overall structure of the study. We aggregate analyses that use the same workflow template into a combined *layer* node. Although the analyses use the same workflow template, they may vary in the number of incoming and outgoing files and the number of contained analysis input groups. We describe these variations—called *layer delta*—in the analyses by computing the similarity of all analyses within one layer based on the number of incoming and outgoing files and the number of contained analysis input groups. To make them comparable, we calculate the difference of each analysis to the analysis with the earliest execution time (within the same layer). Figure 3 illustrates the result of the motif-based aggregation from analysis nodes (AL2) to the layer aggregation level (AL3), resulting in an even higher compression.

**Figure 3:**
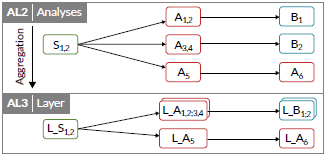
AL2 corresponds to AL2 in Figure 2(b). The additional aggregation level AL3 groups similar analyses into layers using motif-based aggregation.

### 5.2 User-Interest Driven Expansion

Based on a modular degree-of-interest (DoI), we expand regions that are of particular interest to the user. Our interest-driven expansion corresponds to an unbalanced drill-down [EF10] and enables the user to investigate nodes on lower aggregation levels, while keeping the overall graph as a context to analyze the driving changes in the development of a study (e.g., workflow executions, recurring executions, and changes).

Each node has a DoI value assigned that reflects the current level of interest and controls its aggregation. We identified and implemented five components that contribute to the DoI of a node, grouped into user actions and analysis attributes. The components *filter* part of the facet-browsing interface (see Section 5.4), *highlight* when hovering over a node, and *selected* when clicking a node are driven by the user, whereas *workflow execution time* and *layer delta* are based on node attributes (see Figure 4). Each component consists of a weight, configured in a user interface and the value provided by the node itself. The modular Dol function integrates all Dol components into a single Dol value using the following equation:

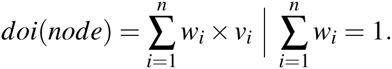

The DoI value for a given node is equal to the sum of all component weights times the component values. The applied weights can be defined freely by the user, where *n* is the total number of DoI components, *w* are the weights of the components, which sum up to one, and *v*_*i*_ is the attribute value. The values for the user actions are binary, meaning the corresponding DoI component value is set to 1 (active = highlighted, selected, or passing a filter) or 0 (inactive). For the values of analysis attributes we use a continuous and normalized scale ranging from 0 to 1.

**Figure 4:**
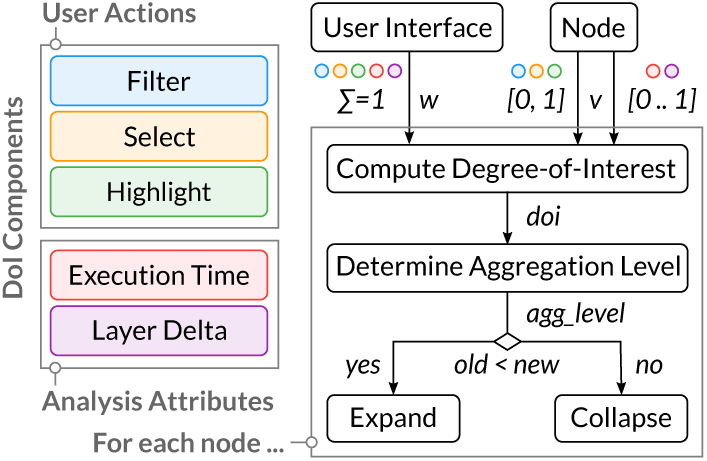
The DoI value for each node incorporates the user-driven weight w and value v for all DoI components. Based on that value the aggregation level of the node is selected, which determines the compression or expansion of the node.

In the next step, the DoI value drives the selection of the aggregation level of the node (see Figure 4). We partition the DoI range into increasing aggregation levels. When nodes switch aggregation levels due to user interaction, we use animated expand and collapse transitions, preserving the mental map of the user. Note that we can also *extract* analyses (AL2) with a high DoI value from a layer node (AL3) *without expanding* them, as shown in the layer far right in Figure 1. With this extraction technique analysis paths can be hidden that are currently not of interest to the user, while still showing interesting ones attached to the layer.

We visualize all Dol components as a stacked bar chart (see Figure 1(b)), which allows the user to manipulate the weight of each component interactively [SGAS16]. When adjusting the weights, the other components adapt their weights proportionally to ensure a sum of 1. Moreover, we allow the user to work with only a single Dol component by activating or deactivating them individually. Applying a modified Dol function results in an update of the provenance graph representation.

### 5.3 Visual Encoding

The data provenance graph contains a series of node attributes that need to be effectively encoded (Task II). Depending on the aggregation level, we create a different node glyph.

**Node Type** Figure 5(a) illustrates the different node types. Nodes in AL0 are visualized as primitive shapes, e.g, diamonds for tools, squares for raw input data, and circles for files. We visually group the workflow using a bounding box with a semi-transparent background that corresponds to the node of higher aggregation levels. Nodes at the level between AL1 and AL3 are drawn as rectangles including a unique icon representing the aggregation level (e.g., a cogwheel for analysis input groups) and the number of aggregated child nodes. in case of layers (AL3) we add the total number of child nodes to indicate possible expansions. For a better distinction between analyses (AL2) and layers (AL3), we add a dashed outline to the layer bounding box to indicate the aggregation.

**Figure 5:**
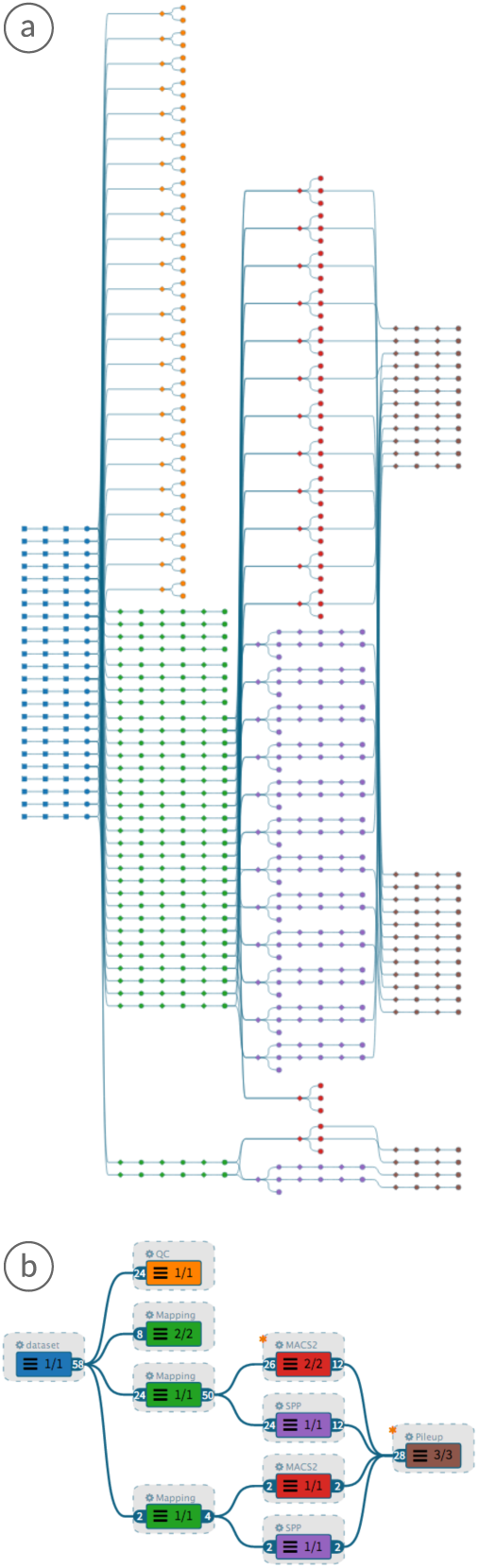
Overview of the data provenance graph of the analysis. Colors represent workflow instances QC (orange), Mapping (green), MACS2 (red), SPP (purple), and Pileup (brown). (a) Fully expanded graph showing tool and file nodes. (b) Graph aggregated to level AL3.

**Age** We vary the brightness of the nodes to encode execution date and time of analyses at all aggregation levels (see Figure 6), addressing Task IV. White represents the earliest and black the most recent analysis execution time. As layers contain analyses that were created at different time steps, we use a black-to-white gradient to encode the distribution of execution times within the layer node.

**Figure 6:**
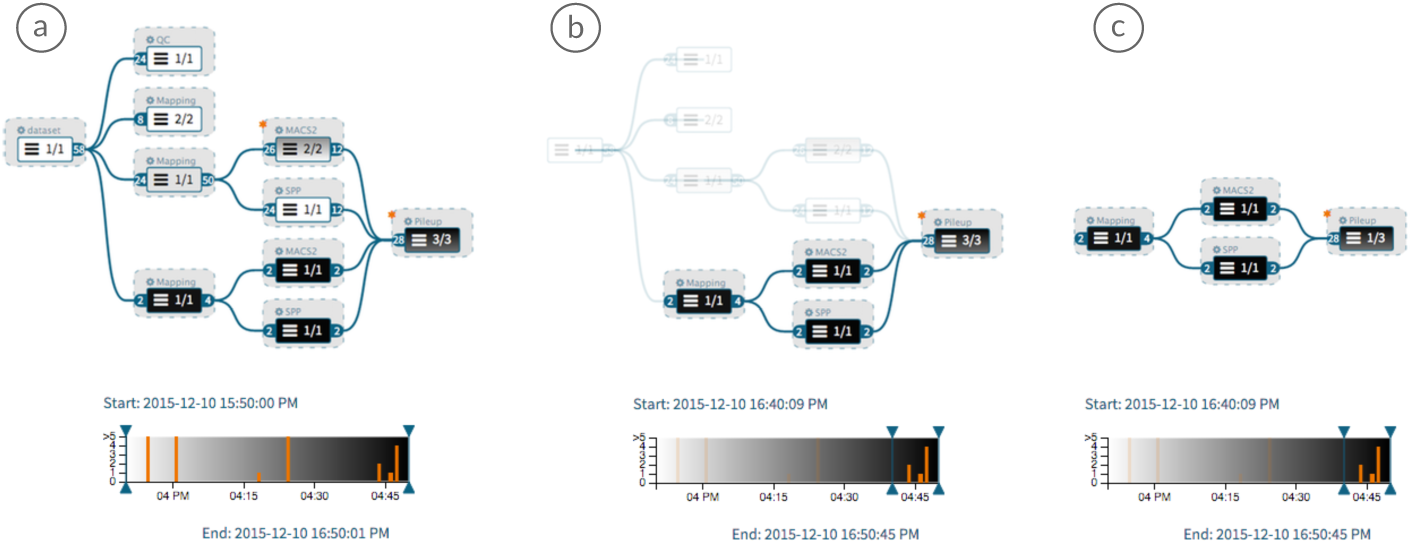
Attribute mapping and time-based filtering. Darker nodes were created more recently. (a) Data provenance graph aggregated to level AL3 with node color representing time. No filtering applied. (b) Time filter applied in blend mode. (c) Time filter applied in hide mode.

**Change** We present the layer delta as defined in Section 5.1 to address Task IV by adding an asterisk—as shown in Figure 1(a)—to the layer node (AL3), in which the similarity value is greater than zero.

**Attribute Information** By applying semantic zooming, we reduce the amount of text labels based on available space. For the aggregated nodes of AL1 to AL3, we show the workflow name next to the bounding box, together with the number of child nodes and the number of incoming and outgoing edges. For workflow instances in AL0, the displayed attribute value (e.g., tissue, factor, or file name) is selectable by the user.

### 5.4 Interaction Techniques

In AVoCADo the user can navigate between the aggregation levels—drill-down (Task III) and roll-up (Task I)—for each node individually or use the buttons in the toolbar (see Figure 2(b)) that directly set all nodes to the selected aggregation level. in addition, the user can search or filter nodes and highlight specific paths to understand causality. We use animated transitions for all operations to provide visual continuity and maintain the mental map of the user.

**Filtering** We provide two user interfaces for filtering nodes by attribute value and time (Task VI). A facet-browsing interface limits the number of nodes based on the contained attribute, such as tissue, drug, or cell type. in addition to the attribute-based filter, analyses can be filtered by analysis execution time using an interactive timeline, as illustrated in Figure 6. The date and time is mapped to the x-axis of the timeline, while the analysis input group count is mapped to the y-axis. The background corresponds to the time gradient (see Section 5.3). The user can move filtering sliders to adjust the time range. Hovering over an analysis highlights the corresponding node in the provenance graph view.

All filter operations result in more available screen space for regions with high interest. Nodes that are affected by a filter can either be shown with reduced opacity or hidden, i.e., removed from the graph, as shown in Figure 6(b) and (c). Hide operations require a recomputation of the whole layout. We apply animations to transition to the new layout.

**Path Highlighting** Enabling the user to investigate the steps that led to an analysis result is an important task (Task V). For each graph node we provide the selection of the predecessors, i.e. the path leading to the selected node, and the successors, i.e. all nodes derived from the selection. The path is highlighted by changing the color of the edges (see Figure 1). Additionally, the user can apply the Dol function to expand all nodes along the highlighted path, while keeping the remaining nodes aggregated.

Note that the primary purpose of our interactive AVOCADO visualization is to enable analysts to *recall* already executed multi-step analysis workflows. Although, the presentation of findings is not a primary goal of AVOCADO, it is indirectly supported via static screenshots that can be shared with others.

## 6 Implementation

AVOCADO is implemented in JavaScript, jQuery, and D3 [BOH11] and handles initialization, layout computation, motif-based compression, and rendering. We use the *Dagre* [pet15] JavaScript library for the computation of layered graph layouts for data provenance graphs with hundreds of nodes. *Dagre* implements the 2-layer crossing minimization [JM97] to reduce the number of edge crossings. Additionally, we compute the order of analysis input groups (AL1) using a barycentric heuristic. We also employ horizontal coordinate assignment [BK02] to balance the speed of computation with layout aesthetics. The layout adapts dynamically to user actions (e.g., filtering, collapse/expand) and DoI changes on a per node basis. Changing the DoI of any node recomputes the weighted sum, resulting in an automatic adjustment of the aggregation level of the corresponding analysis. Our motif-based compression algorithm uses a single breadth-first traversal to discover motifs in the topology-sorted graph, as explained in Section 5.1. Analyses are added in a repeated traversal to layers, based on their preceding motif sequence and the motif itself. With this additional aggregation constraint, we avoid layering analyses solely based on their workflow template without considering provenance in preceding layers. We then calculate and normalize the numeric change metrics of every layer. The assembled data provenance graph is stored in a hierarchical data model where all aggregation levels inherit from a generic node object. Finally, the graph and the filter components (i.e., timeline and DoI view) are rendered in SVG using D3 [BOH11].

The data provenance graph visualization, multiple support views, and the user interface are integrated into the dataset browser of Refinery. Our visualization acquires datasets and their provenance graph over an internal RESTful API in JSON format.

## 7 Usage Scenario

We demonstrate the functionality and effectiveness of the AVOCADO approach by describing how it can be applied in a typical scenario in which an analyst needs to recall, review, and interpret data provenance for a study that was conducted by a collaborator (see Supplementary Video).

### 7.1 Data

Here we are using data provenance information from a *simulated* epigenomics study. Epigenomics is the study of naturally occuring biochemical modifications of the genome that influence gene regulation as well as many other essential cellular processes. The prevalent technology to study these modifications is called ChIP-seq (Chromatin Immuno-Precipitation sequencing) [Par09], which is based on high-throughput sequencing. In our example, several workflows were applied to raw ChIP-seq data in order to identify differences in the distribution of two modifications called *H3K27ac* and *H3K4me3* along the mouse genome in two different tissues (*kidney* and *liver*) after treatment with one of three different drugs (*Alpha*, *Beta*, *Mock*). This results in a total of 12 combinations of experimental factors (2 modifications × 2 tissues × 3 treatments) and for each combination data from two replicates was obtained, resulting in a total of 24 input files for this study.

We applied five different state-of-the-art workflows in this study (see Supplementary Figures S4-S8), implementing a typical ChIP-seq analysis approach as described by Park [Par09]. The specific workflows are (1) *QC*: a quality control workflow to evaluate the quality of the input data. The output is a report. (2) *Mapping*: a workflow that maps the sequencing reads in the input data to the genome sequence. (3) *MACS2* and (4) *SPP*: are peak calling tools that identify locations within the genome to which a larger number of sequencing reads have been mapped than elsewhere in the genome. (5) *Pileup*: This is a workflow that prepares the data for visualization in a genome browser tool.

Due to the computational effort and cost that would be required to execute these workflows on real data, we modified the workflows as follows: (1) We replaced each tool in the workflows with corresponding dummy tools that only pass through any input files that they receive. (2) instead of real data, we used small text files. All tools and workflows (see Figure 7) are available on Github [Ref15c] as well as the meta data and data files used to run the analyses in Refinery [Ref15b].

**Figure 7:**
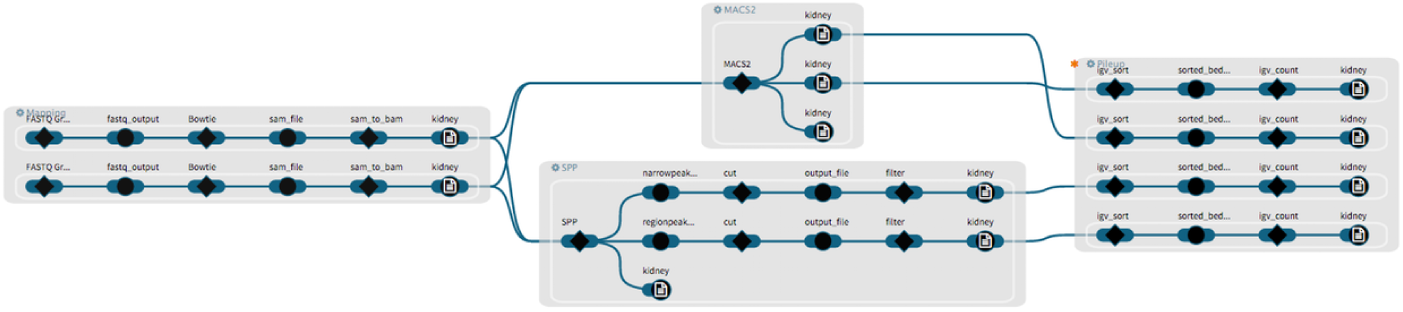
The filtered subgraph from Figure 6(c) is expanded to the workflow instance level (AL0) with the tissue attribute mapped to the output nodes (“kidney”).

Although these analyses were simulated at the tool level, *the simulation has no effect on the size and properties of the data provenance*, which allows us to demonstrate the capabilities of AVoCADo using the resulting graph.

### 7.2 Data Provenance Exploration

As a first step (see Supplementary Video), the analyst wants to gain an understanding of the structure of the study (Task I). While the fully expanded data provenance graph (Figure 5(a)) only provides a hint at the structure of the study, AVOCADO provides a more compact and task-appropriate representation at the highest aggregation level (AL3) (Figure 5(b)). Based on the colors of the aggregate nodes and their labels, the analyst can quickly review which workflows were used and how often. Furthermore, the aggregation of all redundancies in the graph used in level AL3 still provides information about the order in which workflows were chained together. Next, the analyst wants to understand in which order the collaborator ran the analyses and switches the color mapping to the time encoding that maps analysis execution time stamps (Task II) to a unidirectional linear color map (see Figure 6(a)). in this view, the analyst discovers that a part of the analysis was conducted very recently, while the rest was performed much earlier. The analyst then applies the timeline filter (Task VI, see Figure 6(b)) to limit the graph visualization to the most recent analyses and switches filtering into “hide” mode (see Figure 6(c)). Next, the analyst drills down into the most recent set of analyses to the workflow instance level (Task III, see Figure 7). Using the attribute mapper, the analyst maps the tissue attribute onto the output files (Task II) and observes that the most recent analyses were conducted only on *kidney* samples. By hovering over the output nodes of the *MACS2* and *SPP* workflows, the analyst reviews additional attributes of those nodes and observes that both the narrow peak and the region peak files of these tools were compressed with the *Pileup* workflow. The analyst returns to the AL3 level and decides to investigate a layer node that has a change indicator (Task IV) and contains analyses that were executed at different time points (see Figure S1(a)). After drill-down into the analyses (see Figure S1(b) and (c)), the investigator finds that the collaborator re-ran the *MACS2* workflow on a pair of files but that the results were not processed any further. Following this observation, the analyst decides to investigate what the raw data associated with those files is, in order to understand why these files were not processed. By switching the DoI function to always expand selected paths to the highest level of detail, the analyst selects the inputs of the *MACS2* workflow and traces them back to the raw data (Task V, see Figure S2(a) and (b)). Once the raw data files are identified, the analyst then applies the reverse functionality and traces all results generated from one of the raw data files. The analyst observes that this file was processed by both the *MACS2* and *SPP* workflows and that results of both tools were also processed with the *Pileup* workflow (see Figure S3).

This usage scenario illustrates how AVOCADO can be applied in a typical analysis session to address the tasks that we defined with domain experts and based on the literature (see Section 3). In order to evaluate the AVOCADO approach for a wider spectrum of scenarios and to refine it to also handle more extreme cases, a comprehensive user study will have to be conducted as outlined in Section 9.

## 8 Discussion

Graph Layout The choice of layout algorithm is a crucial factor in the perception of a graph visualization. We use a grid-based approach as described in Section 5. The largest node (e.g., an expanded workflow) defines—similar to a spreadsheet—the width for the remaining cells in the same column, which results in long edges to nodes in adjacent columns. Other layout approaches might create a more compact layout (e.g., [YDG*15]), but must also consider the characteristics of the data provenance graph (see Section 2) and the interactive requirements to ensure fluid transitions when changing hierarchy levels. Another issue is the number of edge crossings, which decreases the readability of the graph. To minimize the number of edge crossings, we re-order the nodes in a post-processing step after creating the initial layout. A further aspect is the layout stability. In AVOCADO we initially compute a stable layout, as explained in Section 6. However, interactions such as drill-down/roll-up and DoI expansion change the size of one or multiple nodes and therefore trigger a re-computation of the whole layout. This is neither resource nor time efficient. In future work, this can be improved by recomputing only parts of the graph that have changed together with their neighboring context to make space in the layout.

**Scalability** The AVOCADO approach relies on the hierarchical structure of the provenance graph to achieve compression across the three aggregation levels AL1 through AL3. in order to scale, AVOCADO requires that each level has fewer nodes than the level below, which is generally the case for biomedical workflows. For each aggregation level the same scalability considerations as for general node link diagrams apply. The current implementation performs well on typical biomedical workflows that have around 1,000 nodes on AL0, but technical limitations lead to decreased response times when the provenance graph grows beyond that size.

**Degree-of-Interest Function** The effect of a multi-component Dol value can be hard to interpret. in practice, we observe that users tend to distribute the weights equally or select only a single component at a time. A set of predefined configurations for different tasks (auto-expand selected paths, fully expand clicked nodes, etc.) would simplify the Dol interface but requires further user testing.

## 9 Conclusion and Future Work

In this paper we presented AVOCADO—an approach for visualizing workflow-derived data provenance graphs. We visualize the multi-attribute time-dependent graph as a node-link diagram. To reduce the size of the graph, we apply a combination of hierarchical and motif-based graph aggregation. We interactively expand parts of the graph based on a modular Dol function that is based on graph attributes and user-driven actions. in a usage scenario we demonstrate how our technique can help analysts to gain a deeper understanding of complex multi-step analyses.

In the future we plan to conduct a user study to investigate how to balance multiple competing aggregation strategies effectively. in addition, we want to refine our technique to support a wide range of use cases and scale the visualization to thousands of nodes.

A deeper integration of AVOCADO into tools that capture and manage data provenance information, such as the *Refinery Platform*, will be a step closer towards *actionable provenance*. This will enable analysts to replicate previous analyses results and also to relaunch particular analyses with a new or a different dataset or parameters directly from the visualization. Analysts will not only be able to explore provenance information but also create new data provenance in a single visual interface.

## 10 Acknowledgements

We are grateful to Sam Gratzl for input on the early design and the implementation of AVOCADO and the Refinery Platform team (Peter J Park, Shannan Ho Sui, Win Hide, Ilya Sytchev, Jennifer Marx, Scott Ouellette, Fritz Lekschas) for their help with the task definitions and the integration of AVOCADO. This work was funded by the Austrian Research Promotion Agency (FFG 840232), the Austrian Science Fund (FWF P27975-NBL), the State of Upper Austria (FFG 851460), US National Institutes of Health (R00 HG007583), and the Harvard Stem Cell Institute.

